# Pupil diameter tracked during motor adaptation in humans

**DOI:** 10.1101/2021.04.03.438075

**Authors:** Atsushi Yokoi, Jeffrey Weiler

**Author notes:** Correspondence: Atsushi Yokoi.

## Abstract

Pupil diameter, under constant illumination, is known to reflect individuals’ internal states, such as surprise about observation and environmental uncertainty. Despite the growing use of pupillometry in cognitive learning studies as an additional measure for examining internal states, few studies have used pupillometry in human motor learning studies. Here we provide the first detailed characterization of pupil diameter changes in a short-term reach adaptation paradigm. We measured pupil changes in 91 human participants while they adapted to abrupt, gradual, or switching force field conditions. Sudden increases in movement error caused by the introduction/reversal of the force field resulted in strong phasic pupil dilation during movement accompanied by a transient increase in tonic pre-movement baseline pupil diameter in subsequent trials. In contrast, clear changes in pupil responses were absent when the force field was gradually introduced, indicating that error drove the changes in pupil responses. Nevertheless, we found an association between baseline pupil diameter and awareness of the gradually-introduced perturbation assessed post-experimentally. In all experiments, we also found a strong co-occurrence of larger baseline pupil diameter and slower reaction and movement time after each set break. Interestingly, error-induced pupil responses gradually became insensitive after experiencing multiple reversals. Collectively, these results suggest that tonic baseline pupil diameter reflects one’s belief about environmental uncertainty, whereas phasic pupil dilation during movement reflects surprise about a sensory outcome (i.e., movement error), and both effects are modulated by novelty. Our results provide a new approach for non-verbally assessing participants’ internal states during motor learning.

## Introduction

Motor learning, as a process of correcting movements to achieve a goal, is a complex mixture of multiple processes (1–3). For instance, riding a bicycle initially requires substantial effort to control the handlebars and pedals while balancing, but these conscious efforts are eventually taken over by more automatic and implicit control processes. Studies using arm reaching posit that motor adaptation to a novel dynamic/kinematic environment consists of multiple processes, which are often contrasted as conscious/explicit and automatic/implicit components (4–6), each separately depending on prefrontal and cerebellar function (7, 2). Indeed, human functional imaging studies have revealed that during the acquisition of new motor skills, prefrontal and hippocampal areas are particularly active in the earliest stage of learning. In later stages, the motor and parietal areas, as well as subcortical regions, become more active (8–11). However, the precise cognitive processes represented by early prefrontal/hippocampal activation and the ways in which these processes evolve at finer temporal scales remain unclear.

Pupil diameter under constant illumination is known to reflect a variety of internal cognitive states of individuals performing cognitive or simple motor tasks. Task-evoked changes reflected in phasic pupil dilation have been associated with (unsigned) prediction error or surprise about observations (12–18), and mental/physical effort (19–21). Relatively slow changes in tonic baseline diameter have often been associated with arousal/vigilance (22–27), the tendency of the exploration/exploitation trade-off (28, 29), and more recently, subjective uncertainty about the environment (12–17, 30). Although these constructs have different names, they are closely related to each other in terms of their dynamics. For instance, a surprising observation (i.e., a large deviation from an expectation) may imply a change in the environment leading to a (transient) increase in subjective environmental uncertainty. In an uncertain situation, an animal may need to make more mental/physical effort to find a better solution (i.e., exploration) which may recruit increased arousal/vigilance. As the animal adapts to the new environment, surprise about observations and other variables gradually returns to the average level. Thus, surprise and uncertainty appear to be essential for interpreting pupil responses in various tasks.

More importantly, surprise and uncertainty are believed to play a crucial role in learning behavior by dynamically adjusting learning rate, especially in reward-based learning, including conditioning (31) and choice decision-making (15, 32–34). Thus, these constructs could similarly affect motor learning, such as driving explicit processes to explore potential reach plans (5) or changing sensitivity to errors (35). However, there have been surprisingly few attempts to assess the trial-by-trial changes in pupil diameter during human motor learning. In the current study, to characterize pupil responses and the types of information they reflect in a commonly-used motor learning paradigm, we ran a series of experiments in which we simultaneously tracked pupil diameter during short-term force field adaptation in a reaching paradigm (36). In our experiments, we recruited 91 participants who performed reaching movements under the presence of a velocity-dependent force field that was introduced either abruptly (n = 28), abruptly and then reversed multiple times (n = 29), or gradually (n = 34). Our data suggest that the tonic baseline pupil diameter reflects participants’ belief about environmental uncertainty, whereas the phasic pupil dilation during movement reflects surprise about a sensory outcome (i.e., movement error), and both are modulated by the novelty of perturbation. The current study thus provides an interesting new approach for non-verbally assessing participants’ cognitive states during motor learning.

## Materials and Methods

### Participants

We recruited 28, 30, and 35 right-handed participants with no history of neurological disorders for Experiments 1, 2, and 3, respectively (58 males, 35 females; age: 19–37 years). Participants provided written informed consent before taking part in the study. All of the experimental protocols were approved by the ethical committees of the University of Western Ontario (Experiment 1), Osaka University (Experiment 2), and the Center for Information and Neural Networks (Experiment 3). The sample sizes were selected on the basis of previous studies focusing on pupil size as a main variable of interest (15, 37, 12, 38).

### Experimental settings

#### General settings

Participants were instructed to perform a straight center-out reaching movement from a starting position to a goal target in a two-dimensional plane while holding the handle of a robotic manipulandum. The starting position (white circle, 1.6-cm diameter), the target (gray circle, 1.6-cm diameter), and participants’ current hand position (white dot cursor, 0.5-cm diameter) were displayed on liquid-crystal display (LCD) monitors. Throughout the task, we monitored the participants’ eye gaze and pupil diameter using eye trackers (EyeLink 1000, SR Research Ltd., Ontario, Canada). To reduce the impact of unnecessary eye movements on pupillometry measurement, participants were required to maintain fixation on the center of the goal target. Before the main experimental session, participants underwent a familiarization session with continuous visual feedback of their current hand position (i.e., the cursor). In the main experimental sessions, online visual feedback of the cursor during movement was removed and only terminal feedback was provided at movement offset. For both the familiarization and main session, participants were instructed to aim directly at the target. The colors used for the start position (gray), target (gray/green), background disk (pale blue), and endpoint feedback (magenta) were adjusted to ensure that they were approximately isoluminant (details provided in Table 1).

**Table 1.** Luminance value measurements for display colorss.

|  | Exp 1 | Exp 2, 3 |
| --- | --- | --- |
| Black | 0.6 | 0.4 |
| White | 178 | 249 |
| Gray | 80 | 107 |
| Green | 80 | 106 |
| Magenta | 80 | 108 |
| Pale blue | 80 | 105 |

Measured by CA-100 Plus Color Analyzer (Konica Minolta Sensing Americas, Inc., NJ, USA) for Exp 1 and by SR-3AR Spectroradiometer (Topcon Technohouse, Corp., Tokyo, Japan) for Exp 2, 3. Units are in (cd/m<sup>2</sup>).

At the start of each trial, the robot’s handle automatically moved the participants’ hand into the starting position. During a trial, participants maintained the cursor at the start location for 1 s while maintaining fixation on the center of the target. Following this period, we measured participants’ pupil diameter for a variable duration (3–11 s. The start position then changed to green, informing the participants to initiate reaching. Movement was defined as the period at which the hand movement velocity was above a threshold (3.5 cm/s). At the completion of a reaching movement, endpoint feedback was provided by a magenta cursor (0.5 cm diameter) for 1,000 ms.

We introduced a velocity-dependent curl force field (39) to establish the relationship between the pupil response and the motor adaptation. The force field was applied according to the following equation:

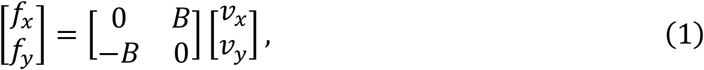

where *f*_*x*_and *f*_*y*_ are the force applied to the handle (N), and *v*_*x*_and *v*_*y*_ are the velocity of the handle (m/s) for the *x*-and *y*-directions, respectively. For the clockwise (CW) force field, the viscosity coefficient B (N/[ms^-1^]) had positive values, and for the counter-clockwise (CCW) field, B had negative values. To quantify adaptation to the force field, we occasionally introduced “channel” trials, in which the handle motion was constrained to a straight path between the home position and the target by a simulated damper and spring (40), to measure the force applied to the channel.

To make inter-participant comparisons of pupil diameter interpretable (e.g., for Exp 3), we additionally measured reference physiological response amplitudes of pupil diameter within each participant while eliciting a pupillary light reflex by changing the background color of the display from light blue to white (higher luminance) or black (lower luminance) (Table 1). These measured pupil limits were used for the within-individual normalization of pupil diameter data (Fig. 1B).

**Figure 1.**
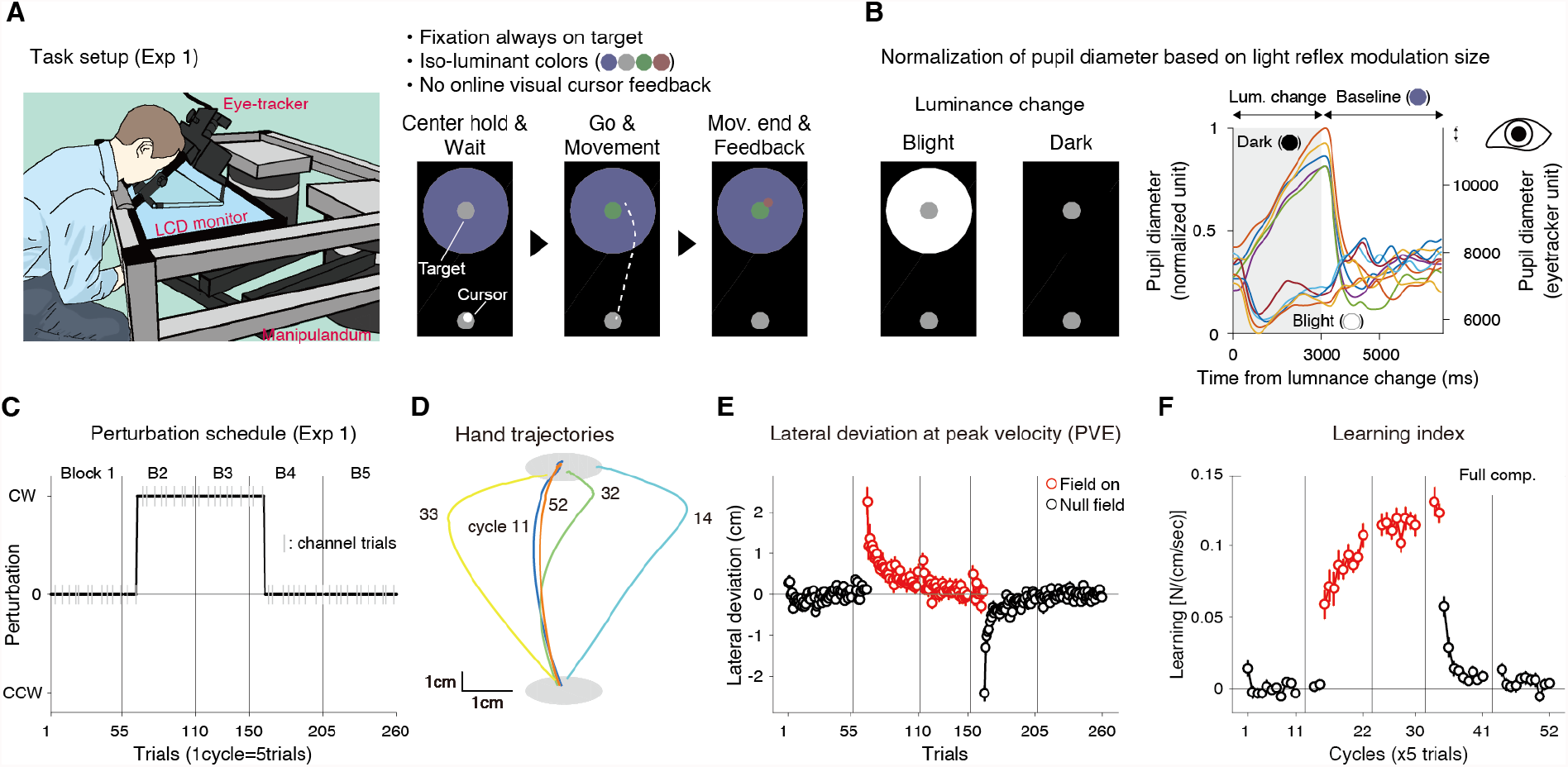
Pupillometry during force field motor adaptation (Experiment 1). **(A)** Left: Experimental setup used in Exp 1 (n = 28). Right: Schematics for a single trial. Participants were required to fixate on the center of the target. **(B)** Pupil diameter data were individually normalized using modulation size of pupil light reflex induced by blight/dark background color change (left). Example time courses of pupillary responses in one participant are shown on the right. Different colors indicate individual trials. **(C)** The perturbation schedule for Exp 1. The manipulandum applied the clockwise (CW) force field unexpectedly for the participants in the second block. The gray vertical lines indicate set breaks. The short gray vertical lines represent the force channel trial where learning was quantified. **(D)** Average hand trajectories of participants who learned with the CW force field. The cycle numbers (11, 14, 32, 33, and 52) correspond to baseline, early perturbation, late perturbation, early washout, and late washout trials. **(E)** Mean lateral hand deviation at the peak tangential handle velocity (PVE). Positive values correspond to rightward deviation. **(F)** Mean learning index measured in the channel trials (once in a cycle). Learning index was the lateral force to the channel at the time of peak velocity divided by the peak velocity (i.e., viscosity). The red dots represent data for perturbation trials (E, F). Error bars correspond to the standard error of the mean (s.e.m). across the participants (E, F).

#### Experiment 1

Experiment 1 was conducted at the Brain and Mind Institute, University of Western Ontario (Ontario, Canada). The self-reported right-handed participants (n = 28; 13 males, 15 females; age: 24.3 ± 4.5) sat on a height-adjustable chair and held the handle of a robotic manipulandum (1,000 Hz control rate) (41). The position of the handle was represented as a cursor (white dot) on an LCD monitor (60 Hz update rate) placed directly above the handle to prevent the participant from seeing their hand. The starting position (white circle), goal target (gray circle), and background (light blue circle, 15 cm diameter) to prevent sharp luminance changes around the target were also presented on the display throughout the task (Fig. 1A). The eye tracker was mounted on the display to monitor participants’ eye gaze and pupil diameter (Fig. 1A). The approximate distance between participants’ eyes and the center of the monitor was 16 cm.

An experimental session consisted of five blocks of trials (59 trials per block, except for the fourth block, which consisted of 44 trials). There were short breaks (up to 1 min) inserted between the blocks. On each block, the first and the last two trials were used to measure the simple pupil light reflex of the participants. In each of these “light reflex” trials, the LCD screen was suddenly turned either black or white for 2,000 ms. The pupil response was measured during and up to 3,000 ms after the termination of the black/white color stimulus. The response strengths (i.e., trough for constriction, and peak for dilation) were averaged over the blocks and used to normalize the individual pupil diameter during the main task. Following the initial “light reflex” trials, participants performed 55 trials of the center-out reaching task. On each trial, after confirming stable eye fixation (1,000 ms of fixation without any blink) on the target and the cursor staying within the home position, the goal target turned green after a variable delay (1,000–1,500 ms) cueing the reaching movement. To prevent possible predictive/reflexive eye movement and/or pupil dilation because of a moving cursor, visual feedback of the cursor was removed during movement and participants were instructed to maintain fixation on the center of the target. Terminal end-point feedback was provided with a near-isoluminant magenta cursor for 1,000 ms when the cursor speed was less than 1 cm/s. After the feedback period, the manipulandum handle automatically returned to the home position and the next trial started.

Experiment 1 employed a typical Null-Force-Null paradigm. The force field was introduced at the 11th trial of the second block and removed at the 11th trial of the fourth block (Fig. 1C). Participants were not informed regarding the trial on which the perturbation would be introduced or removed in advance. Twenty participants were presented with the CW force field (*B* = 0.15 in Eq. [1]) and the remaining eight participants were presented with the CCW force field (*B* = −0.15). To quantify adaptation to the force field, we interleaved the channel trials in 20% of the trials. The channel stiffness and viscosity were 7,000 (N/m) and 30 (N/[ms^-1^]), respectively.

#### Experiments 2 and 3

Experiments 2 and 3 were conducted at the Osaka University/Center for Information and Neural Networks (Suita, Japan). Participants sat on a height-adjustable chair and held the handle of a robotic manipulandum (Phantom Premium HF 1.5, 3D Systems, Inc., USA). The force field strength was set to *B* = 0.12 (CW) or −0.12 (CCW). The channel stiffness and viscosity were 2,500 (N/m) and 25 (N/[ms^-1^]), respectively. The position of the handle was visually displayed on a vertically placed LCD monitor (60 Hz update rate) as a cursor (white dot). A starting position (gray circle), a goal target (gray disk), and a background disk (light blue disk, 21.6 cm diameter) to prevent sharp luminance changes around the target were also presented on the display throughout the task (Fig. 3A). The colors (gray, green, light blue, and magenta) were adjusted to be approximately isoluminant (Table 1). A desktop eye tracker (Eyelink 1000, SR Research, Ontario, Canada) was placed under the LCD display to monitor participants’ eye gaze and pupil diameter (Fig. 3A). The approximate distance between participants’ eyes and the center of the monitor was 35 cm for Exp 2 and 44 cm for Exp 3. We assessed participants’ handedness using a Japanese-translated version of the FLANDERS handedness questionnaire (42, 43) which ranges from +10 (perfect right-hander) to −10 (perfect left-hander).

#### Experiment 2

In Exp 2, 30 right-handed individuals participated. Participants were divided into three sub-groups (10 participants each). One participant assigned in Exp 2C did not complete the whole experimental session because of frequent blinks during an experimental trial. As a result, the data from 29 participants (24 males, five females; age: 22.3 ± 2.7; handedness score: 9.9 ± 0.3) were analyzed (10 for Exp. 2A, 10 for Exp. 2B, and nine for Exp. 2C). All three experiments consisted of five blocks of 50 reaching trials with short breaks (∼1 min) between the blocks. Similar to Exp 1, in each block we added two trials of light-reflex measurements before and after the reaching trials. The perturbation schedules were characterized with sudden reversals in force field direction, as summarized in Figure 3B. For Exp 2A, CW force was applied on trials = {61:89, 101:129, 151:159, 170:179, 190:209, 220:229}, and CCW force was applied on trials = {90:100, 130:150, 160:169, 180:189, 210:219, 230:239}. For Exp 2B, CW force was applied on trials = {70:79, 90:109, 120:129, 140:189, 201:229}, and CCW force was applied on trials = {61:69, 80:89, 110:119, 130:139, 190:200, 230:239}. For Exp 2C, CW force was applied on trials = {61:159}, and CCW force was applied on trials = {160:170}. For all groups (Exp 2A through 2C), channel trials were randomly interspersed in 20% of trials, except that for Exp 2C, channel trials were repeated from the 171st trial to the 250th (last) trial.

#### Experiment 3

In Exp 3, 35 right-handed individuals participated. Of these participants, the data from one participant were excluded from the analysis because of a technical failure in collecting pupil calibration data. Thus, the data from the remaining 34 participants were analyzed (20 males and 14 females; age: 22.2 ± 1.6 years; handedness score: 9.9 ± 0.9). In this experiment, the force field was gradually introduced over the course of seven blocks (50 trials in each block, except for the last block, which consisted of 70 trials). The force field was first introduced at the 16th trial in the second block and incrementally increased by 5% of the full-strength (*B* = 0.12) after every 11 trials, until it reached the full-strength (Fig. 5A). Channel trials were randomly interspersed in 20% of trials, except for block seven, in which channel trials were repeated from the 321st trial to the 370^th^ (last) trial. Nineteen participants adapted to the CW force field, and the rest of the participants adapted to the CCW force field. The force and kinematic data were sign-flipped for the CCW participants. After the experiment, 32 participants in Exp 3 answered a questionnaire asking if they were aware of force perturbation. The questions included (a) whether they felt any load in any phase of the experiment (yes/no); (b) which direction of load they felt (1: forward; 2: backward; 3: rightward; 4: leftward; 5: other); and (c) in which of the seven blocks they felt the load (1–7, multiple answers allowed). In the current experiment, the data for question (c) were analyzed.

## Data analysis

The data were sampled at 200 Hz. All analyses were conducted using custom-written codes with MATLAB R2015b (Mathworks, Natick, MA, USA).

For all of the experiments, the data from the manipulanda (x-y positions, x-y velocities, and x-y command forces for the manipulandum handle) were smoothed with a Gaussian kernel of 35 ms full width half maximum (FWHM). Reach onset was defined by the time at which the tangential hand velocity first exceeded 10% of its peak value. Reach offset was defined as the time at which the tangential hand velocity first dropped below the threshold determined for reach onset of that trial. To assess movement errors on each reach, we computed peak velocity error (PVE) and endpoint error (EPE). PVE was defined as perpendicular hand displacement at the time of peak tangential hand velocity with respect to a straight line from the home position to the target, and EPE was defined as the distance between the hand and the target positions at movement offset. As an index of motor learning, we used the x-force values at the time of peak tangential hand velocity divided by the peak velocity during the channel trials (44). For Exp 1 and 3, the PVE and learning index were sign-flipped for participants who learned with the CCW force field before taking the group-average.

For all of the experiments, the eye position and pupil data (x-y point-of-gaze position and pupil diameter) were analyzed in the following way. First, we discarded 100 ms of point-of-gaze position and pupil data prior to a blink event and 150 ms of point-of-gaze position and pupil data after a blink event. We then interpolated over the discarded data using a piecewise cubic Hermite interpolating polynomial (a Matlab *interp1*.*m* function with “pchip” option). After the interpolation, the eye position data were smoothed with Gaussian kernel with 35 ms FWHM. For the detection of saccade events, the unsmoothed eye position data were also smoothed with a second-order Savitzky-Golay filter with a frame length of 55 ms (*sgolayfilt*.*m* function). The eye velocities were then calculated by the numerical derivative of the eye positions (*diff*.*m* function). We employed 30 °/s as the velocity threshold for saccade detection. Because the eye positions were defined in the display coordinates, we transformed them to visual angle coordinates before the saccade detection.

To remove high-frequency noise, the pupil data were smoothed with a Gaussian kernel with 235 ms FWHM. Importantly, we individually normalized the pupil diameter data relative to the minima and maxima of the pupil diameter data measured during the light reflex trials. We also calculated the pupil dilation velocity through the numerical derivative (*diff*.*m* function) of the normalized pupil diameter data. The trial-by-trial summaries of these pupil-related variables were defined as follows. The baseline pupil diameter was defined as the average pupil diameter during the waiting period before the onset of the go cue. The mean pupil dilation velocity at each trial was defined as the average of pupil dilation velocity during the period from 300 to 700 ms from movement onset. These periods were defined in a post hoc manner to maximally reflect the effect of experimental manipulation (i.e., force perturbation) and roughly corresponded to the periods of *p* < 0.001 (uncorrected) for the comparison between the baseline vs. the first five perturbed trials (e.g., Fig. 2B). Pupil dilation velocity data for trials in which a saccade was detected during the movement period were excluded from the analysis. The percentages of excluded trials were 16.5% ± 12.9% (Exp 1), 15.5% ± 15.2% (Exp 2), and 10.5% ± 8.8% (Exp 3).

**Figure 2.**
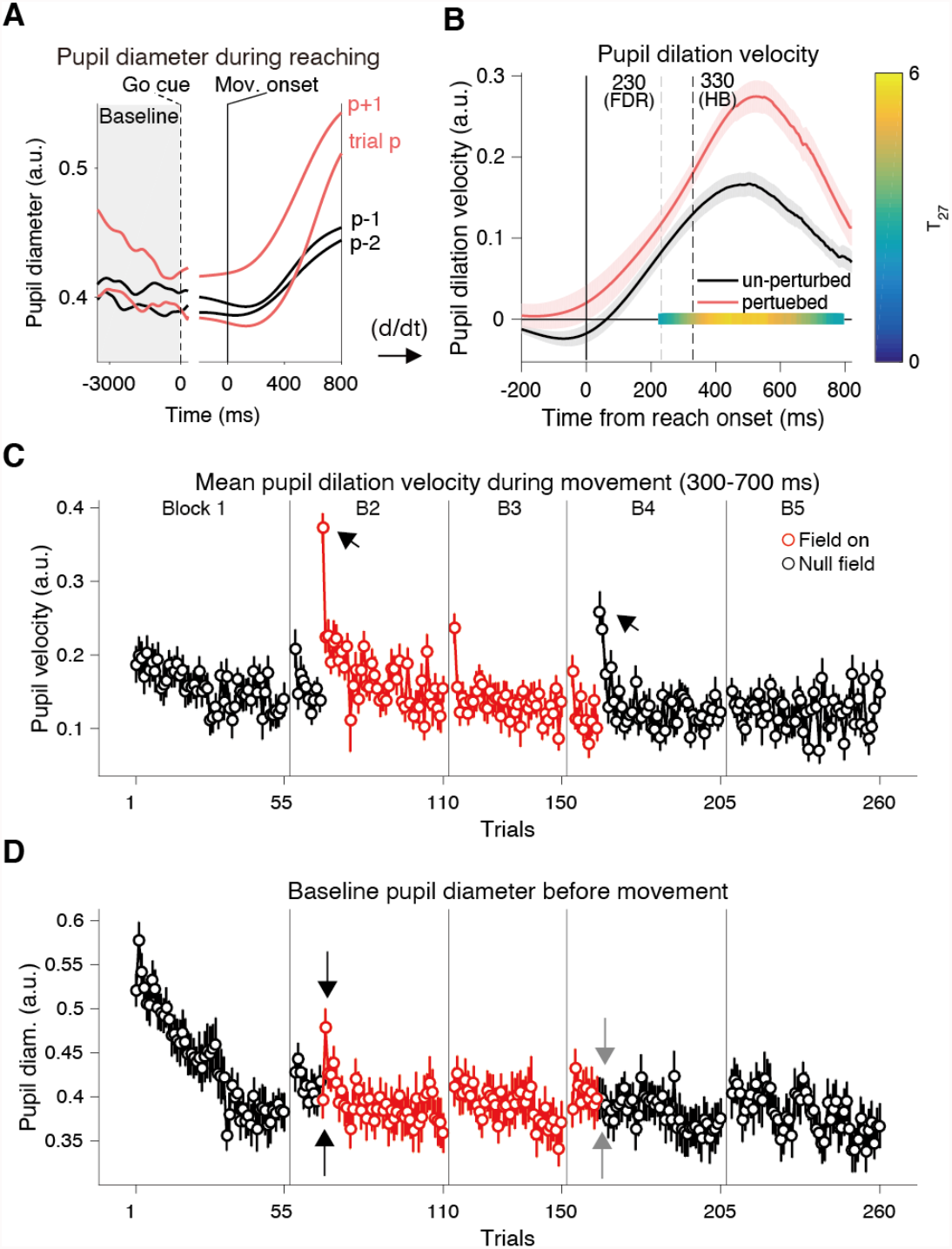
Pupil responses to abruptly introduced force field (Experiment 1). **(A)** Averaged pupil diameter time series across participants during Exp 1, aligned either to the go cue or reach onset. In general, the pupil showed different baseline values and dilation during reaching. On a sudden perturbation trial (indicated as trial p), the pupil was strongly dilated (see also arrows in panel C). On the following trial (trial p+1), pupil diameter showed a higher baseline value (see also arrows in panel D). **(B)** The phasic pupil response to the force perturbation; the first time-derivative (= velocity) of the pupil diameter time series (A) averaged over the trials before the force field was introduced (unperturbed; black trace) or the first five perturbation trials at the second block (perturbed; red trace). The areas around the traces represent the s.e.m. across the participants. The vertical dashed lines indicate the first time point of significant difference assessed by running paired *t*-tests (two-sided) corrected by either the Holm-Bonferroni method (HB; *p* < 0.05) or the Benjamini-Hochberg method controlling for false-discovery rate (FDR; *q* < 0.05). A significant (*p* < 0.05) cluster assessed by cluster-based permutation test is also shown on the y = 0 line, with color-mapping representing *t*-values for comparison (T27; the color bar on the right). **(C)** Trial-by-trial changes in the pupil dilation velocity averaged between 300 to 700 ms from movement onset. **(D)** Trial-by-trial changes in the baseline pupil diameter (mean pupil diameter during the hold and wait period; see panel A). The red circles represent data for perturbation trials (C, D). Error bars correspond to the s.e.m. across participants (C, D). The arrows in (C) indicate the sharp increases in mean pupil dilation velocity at the first introduction/removal of the force field. The arrows in (D) indicate the first two perturbation trials (upward: the first trial, downward: the second trial), and the first two trials since force removal (upward: the first trial, downward: the second trial).

### Statistical analysis

*Time-series comparison:* We assessed changes in the pupil velocity time series between unperturbed (average of the first block data) vs. perturbed (average over the first five perturbed trials) trials using group-wise two-sided paired *t*-tests applied at each time frame. Uncorrected *p*-values were reported. Correction for multiple comparisons (# of time frames since movement onset) was applied using the Holm-Bonferroni method (45) with a family-wise error rate of *p* = 0.05 and the Benjamini-Hochberg method (46) with a false-discovery rate of *q* = 0.05. For a complementary analysis, we also employed the cluster-mass permutation test (47) implemented as ‘*permutest*.*m’* for MATLAB (48) with a significance level of *p* = 0.05 using a cluster-thresholding *p*-value of 0.01. However, it should be noted that the cluster-based permutation test does not establish the significance of effect latency (49).

#### Effect of repetitive change points for Exp 2

We assessed the effect of participants’ repetitive experience of the change points by fitting a general linear mixed-effects model (*‘fitlme*.*m’* function) to the individually z-scored response data around the change points (baseline pupil diameter, pupil dilation velocity). All models included a fixed intercept and random intercepts for subjects and groups (Exps 2A, 2B, and 2C) and fixed and random effects of the perturbation types (CW and CCW). As effects of interest, fixed and random effects for the number of change points (#cps) were also included. To account for the trivial effect of error magnitude, we additionally included the absolute error term: trajectory error and endpoint error at the immediately previous trial for the baseline pupil data, and trajectory error at the same trial for the pupil dilation velocity data. The subject, group, and perturbation types were treated as categorical variables. The importance of the random effect of #cp was assessed using the likelihood-ratio test (LRT) between the full model and an alternative model lacking the random slope term. The models were fitted to the data using the restricted maximum likelihood method (ReML) with random starting values. We evaluated the *p*-value of the estimated fixed-effect slope for the #cp term to test whether the effect of change in #cp was statistically significant. The significance level was set to 0.05.

#### Sub-group comparison for Exp 3

The data for 32 participants who went through the post-study questionnaire about perturbation were analyzed. We first split the data into two sub-groups based on the median value of the total score of the post-study questionnaire (i.e., the total number of blocks in which the participant was aware of perturbation, ranging from 0 to 7). The difference in the score (i.e., number of “yes” responses) between each sub-group was assessed using a chi-squared test at each block. We then compared the pupil and behavior data between the sub-groups in the following way. Data were first averaged within each cycle (five-trial bins), and a linear mixed-effects model was fit to the data. The model contained a fixed intercept, a random intercept regarding subjects, a fixed effect for cycle number within each block, random effects for cycle number regarding subject and block, and a fixed interaction between block and sub-group as an effect of interest. A significant difference between the sub-groups was assessed by the significance of this interaction term for each block. The cycle number was treated as a continuous variable, and other variables were treated as categorical. The models were fit to the data with the ReML method with random starting values. We employed the Holm-Bonferroni method (45) for correction for multiple comparisons (i.e., blocks) to maintain the family-wise error rate at *p* = 0.05. For less-stringent correction, we also used the Benjamini-Hochberg method (46) to maintain the false-discovery rate at *q* = 0.05.

#### Effect of set break (block novelty)

To test the effect of starting new blocks avoiding the effect of perturbation, we first selected the blocks in each experiment in which perturbation was either absent or very weak for the initial 10 trials (blocks 1, 2, and 5 for Exp 1, 1 and 2 for Exp 2, and 1–4 for Exp 3) and averaged the data within each trial defined relative to block initiation. We then compared the average of the initial two trials (early) vs. the 6th–10th trials (late) for baseline pupil diameter, pupil dilation, reaction time, and movement time, using a two-sided paired *t-test* for each experiment.

#### Data and code availability

The data and the custom-written Matlab codes used for the analysis will be uploaded to the publicly available server upon publication. Until then, requests should be addressed to the corresponding author.

## Results

### Pupillometry during simple force field adaptation paradigm: Error-driven pupil responses

To obtain initial insights about pupil movement changes during a typical motor adaptation paradigm, we monitored 28 participants’ eye movements while they performed a reaching task with a force field perturbation (Exp 1; Fig. 1A, C). To minimize measurement noise in pupillometry, we instructed the participants to keep their eyes fixated on the target and to refrain from blinking while they performed reaching movements. The visual cursor feedback was occluded during reaching. To minimize brightness-induced changes in pupil size, visual stimuli were isoluminant with respect to the background (Fig. 1A). Participants were instructed to aim directly at the target, and to reach straight to it. Participants showed a typical behavioral signature of force field adaptation. A sudden introduction of the perturbation force disturbed the smooth, relatively straight hand trajectory (cycle 11, Fig. 1D) resulting in a large lateral deviation (cycle 14, Fig. 1D). The lateral deviations rapidly decreased with repeated reaches made under the perturbation (cycle 32, Fig. 1D; Fig. 1E), and the sudden removal of the force perturbation resulted in a large hand deviation toward the opposite direction (cycle 33, Fig. 1C; Fig. 1D), a signature of motor adaptation known as an aftereffect. With further trials, the trajectory (and movement errors) returned to a near-baseline level (cycle 52, Fig. 1D; Fig. 1E). The learning quantified in the force channel trials also showed a typical learning curve (Fig. 1F). How do participants’ pupils respond in this typical motor adaptation situation?

The pupil typically showed phasic dilation during movement in the Null-field trials (Fig. 2A, p-1 and p-2). When the perturbation was unexpectedly applied, the pupil exhibited additional dilation (Fig. 2A, trial p, p+1). Analysis of the time derivative of the pupil dilation (pupil dilation velocity) revealed that this perturbation-evoked pupil dilation started ∼300 ms after movement onset and lasted until ∼700 ms after movement onset (Fig. 2B). The trial-by-trial change in the pupil dilation velocity averaged over this window (we will refer to this as pupil dilation) showed a sharp rise upon the introduction of the force field and gradual decline as the participants adapted to the force field and the movement error decreased (Fig. 2C; Fig. 1E). This indicates that, in the current paradigm, the phasic pupil dilation during reaching did not simply reflect physical effort, as it decreased despite the increase in lateral force exerted by the participants (Fig. 1F; Fig. 2C). The increased pupil dilation was also followed by increased tonic baseline pupil diameter in the following trial (Fig. 2D, black arrows) (*t*_27_ = 5.09, *p* = 2.4 × 10^−5^, two-sided paired *t-*test). Faced with the sudden removal of the force field and the resultant aftereffect, pupil dilation showed a re-increase (Fig. 2C; block 4), whereas the baseline pupil diameter did not show such a re-increase (Fig. 2D, gray arrows) (*t*_27_ = 0.001, *p* = 0.99, two-sided paired *t-*test). It is also noteworthy that the tonic baseline pupil diameter showed higher values at the beginning of a new block (Fig. 2D). We will later provide more detailed analysis regarding these points combined with the results of Exp 2 and 3.

The results described above suggest that the phasic pupil dilation is likely to be modulated by the size of movement error. The correlation between the trial-by-trial change of phasic pupil response and that of absolute movement error for the group-averaged data was highly significant (*r* = 0.64; *p* = 8.97 × 10^−31^). This finding appears to be consistent with recent proposals that task-induced pupil dilation reflects surprise (e.g., unsigned prediction error, or risk prediction error) and the tonic baseline pupil diameter reflects subjective uncertainty about the environment (15, 14). On the one hand, this fits well with the observed change in baseline pupil diameter; an abrupt introduction of the force field strongly implies environmental change leading to a transient increase in subjective uncertainty about the task, which is reflected in the baseline pupil diameter. On the other hand, the sudden removal of the force field did not elicit such changes in the baseline pupil diameter, presumably because the participants were aware that the task has returned to a known “baseline” state, implying an effect of novelty on environmental uncertainty, or simply an effect of fatigue or reduced arousal.

To further investigate the relationships between pupil diameter, error size and environmental change during motor learning, we also monitored the pupil diameter in two additional experiments. In Exp 2, we employed a switching force field schedule, in which the direction of force field was unexpectedly reversed multiple times, inducing overt environmental change and a re-increase in error. In Exp 3, we employed a gradual force field schedule, in which the magnitude of force field was gradually increased, inducing covert environmental change with much smaller errors compared with Exp 1 and 2.

### Pupil responses to a switching force field schedule: Dissociation between error sizes and pupil responses after multiple reversals

In Exp 2, we monitored the pupil responses of another 29 participants while they reached in the presence of force fields with slightly different settings compared with Exp 1, including a light-weight manipulandum and a vertically aligned monitor (Fig. 3A). Exp 2 involved three sub-groups (Exp 2A, B, and C) in which participants experienced different force field schedules (Fig. 3B). In all of the sub-groups, the force field was abruptly introduced on the 11th trial in block 2 (CW for Exp 2A and C, and CCW for Exp 2B), and the direction of the force field was reversed at different timings and frequencies for different sub-groups (Fig. 3B; see Methods for detail). These change points, including the introduction and removal of the force field, induced sudden, unexpected increases in error size (Fig. 3D), indicating a change in task environment. We examined participants’ pupil responses, both phasic and tonic, to these change points.

**Figure 3.**
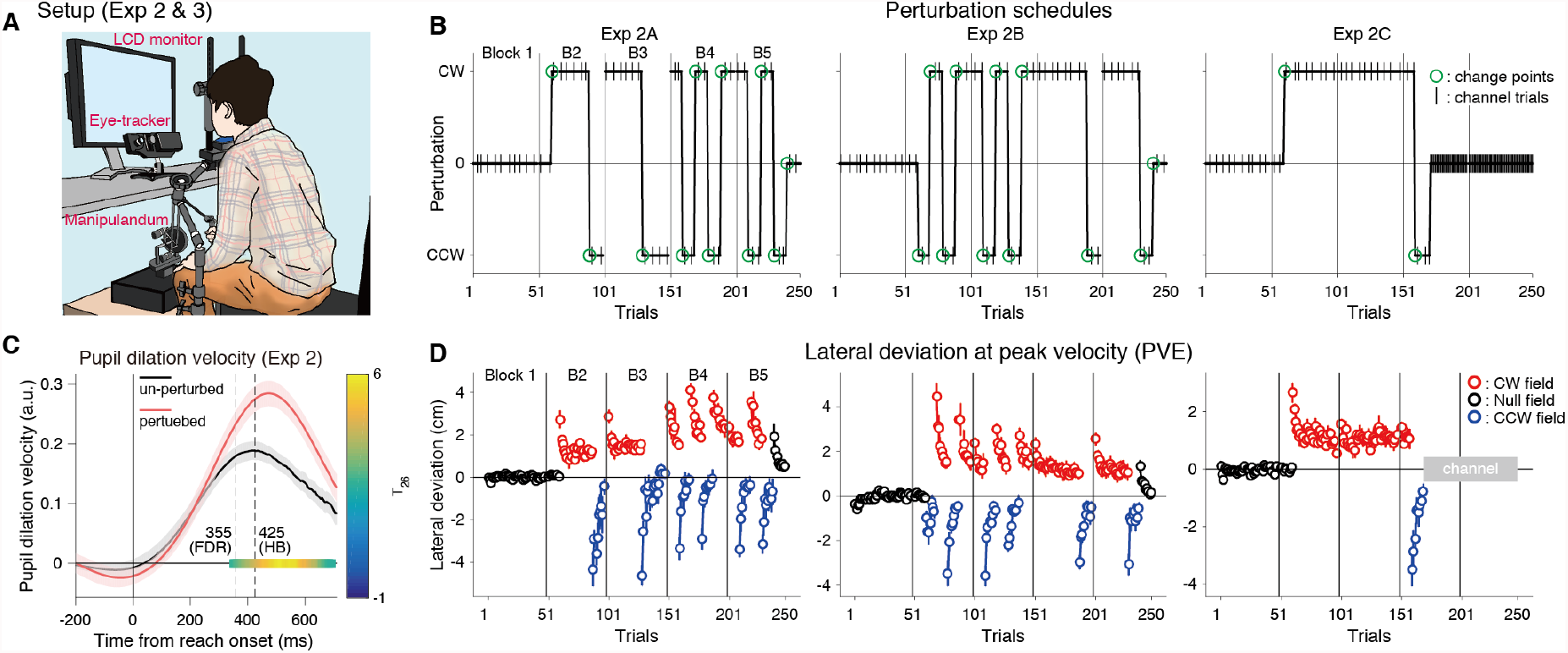
Pupil responses to sudden reversals in the force field. **(A)** Experimental setup for Experiments 2 and 3. **(B)** Perturbation schedules for Exp 2 (Exp 2A [n = 10], 2B [n = 10], and 2C [n = 9]). The green circles indicate the change point trials in which either the magnitude (on/off) or direction (CW/CCW) of the force field changed in the middle of the blocks (changes across the blocks were excluded). The vertical lines indicate set breaks. **(C)** The time course of the pupil dilation velocity averaged over the baseline trials (un-perturbed; black trace) or the first five perturbation trials in the second block (perturbed; red trace), averaged over all sub-groups. The areas around the traces represent s.e.m. across the participants. The vertical dashed lines indicate the first time point of significant difference by running paired *t*-tests (two-sided) corrected with either the Holm-Bonferroni method (HB; *p* < 0.05) or the Benjamini-Hochberg method (FDR; *q* < 0.05). A significant (*p* < 0.05) cluster assessed by cluster-based permutation test is shown as a color map on the y = 0 line representing *t*-values for the comparison (T26 because of missing data; the color bar on the right). (**D**) Trial-by-trial change in the mean lateral hand deviation at the peak tangential handle velocity for the Exp 2A (first column), 2B (second column), and 2C (last column), respectively. The colors of circles (black, red, and blue) indicate the data for baseline, CW field, and CCW field, respectively. Positive values for panel D correspond to rightward deviation. The vertical lines indicate set breaks. Error bars represent s.e.m. across participants. Trials in which the dot color changed indicate the change points.

First, to replicate the basic results of pupil responses in Exp 1, we analyzed the pupil data around the first change point (i.e., the first introduction of force field) aggregating the data from all sub-groups. The time course of perturbation-evoked phasic pupil dilation showed a similar pattern (Fig. 3C) to that in the first experiment (Fig. 2B) with a slight difference in the timing of the effect. Thus, we employed a similar time window for averaging the pupil dilation velocity at each trial (300 to 700 ms). As shown in Figures 4A and C, both the phasic pupil dilation during movement and the tonic baseline pupil diameter responded to the first change point in a similar way to those in Exp 1. How does the pupil respond to the following change points?

**Figure 4.**
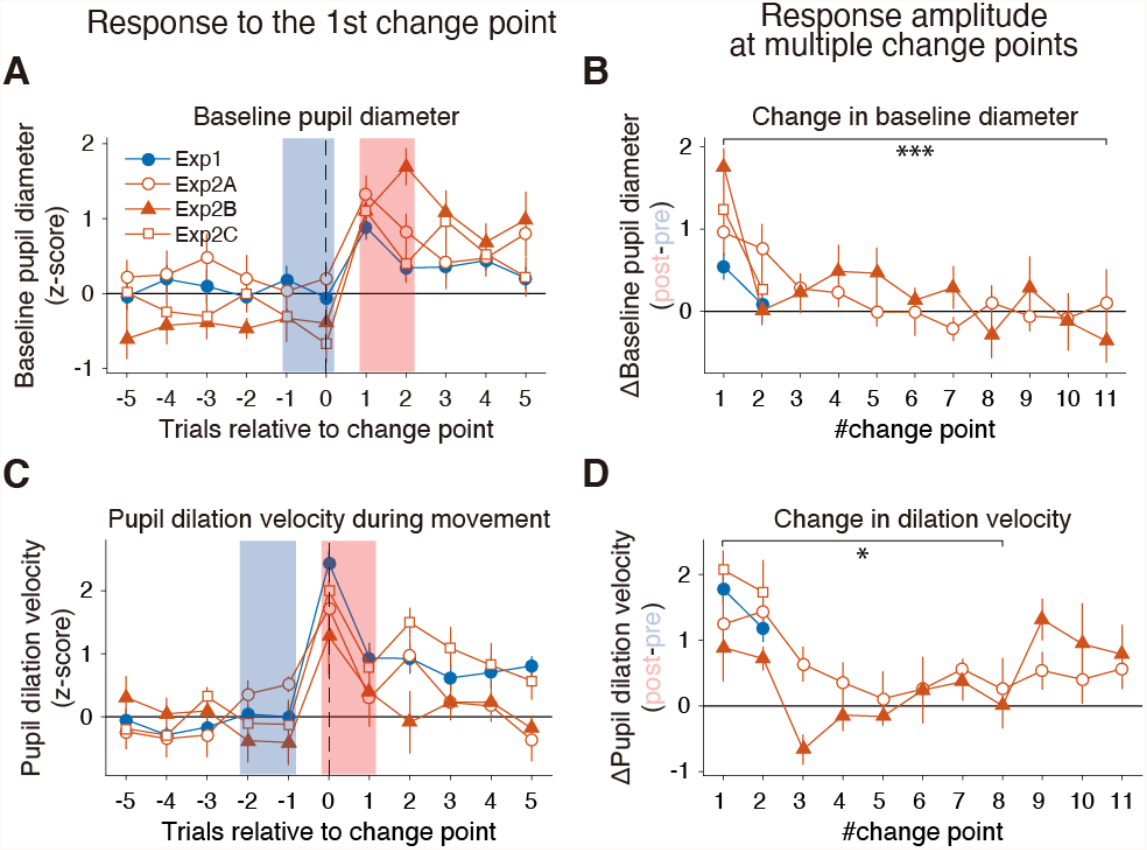
Pupil responses around change points. **(A)** Trial-by-trial change in the z-transformed baseline pupil diameter around the first change point. Pre- and post-change point values are defined by blue and red shading, respectively. **(B)** The amplitude of change point-induced baseline pupil diameter change (difference between post-vs. pre-change point values in A) as a function of the number of change points. **(C)** Trial-by-trial change in the z-transformed mean pupil dilation velocity around the first change points. Similarly, pre-and post-change point values are defined by blue and red shading, respectively. **(D)** The amplitude of change point-induced mean pupil dilation velocity change (difference between post-vs. pre-change point values in C) as a function of the number of change points. The error bars indicate s.e.m. across the participants (Exp 2A [n = 10], 2B [n = 10], and 2C [n = 9]). Asterisks indicate a significant slope for the #change point assessed by the linear mixed-effects model (*: *p* < 0.05; ***: *p* < 0.001; see main text and Methods for more detail).

Intriguingly, despite the sharp re-increase in movement error at the following change points (Fig. 3D), both the pupil dilation and baseline pupil diameter quickly became insensitive to these large errors (Supplementary Figure 1). It should be noted that such a decline in pupil response sensitivity to errors was not reported previously in the context of reinforcement learning (e.g., 14, 15). To quantify the decline in pupil responses, we focused on response amplitudes around each change point. The response amplitude in the baseline pupil diameter and the pupil dilation showed a dramatic decline for the subsequent change points (Fig. 4B, D). Such a decline in pupil response amplitude was also observed in the data in Exp 1 (Fig. 4B, D). A linear mixed-effects model on the baseline pupil diameter confirmed the significantly negative slope on the change point (*t*_234_ = −5.6, *p* = 5.7 × 10^8^) after controlling for the random group/individual factors and the effect of trajectory and endpoint errors at each change point (see Methods). This finding was unexpected because the size of the unsigned error increased almost two-fold at the second change point because of the reversal of force field direction (Fig. 3D). Such reduced sensitivity to error is unlikely to have been caused by fatigue or boredom alone, as participants in Exp 2B already showed a substantial decline in baseline diameter change at the second change point (Fig. 4B) which was still in the second block of the experiment (70th trial). Similarly, the pupil dilation showed decreased sensitivity to errors. Although there was no significant monotonic decrease (*t*_175_ = 0.04, *p* = 0.97, the same linear mixed-effects model without the effect of endpoint error) as seen in the baseline pupil diameter, visual inspection suggested that this was caused by the sudden increase in the phasic pupil response on the 9th change point in Exp 2B (Fig. 4D). The slope up until the 8th change point was significantly negative even after correcting for the error (*t*_116_ = −2.2, *p* = 0.027, linear mixed-effects model), suggesting diminished sensitivity to physical error in the phasic pupil response, similar to the tonic pupil response to the change points. The recovery in response amplitude seen in the pupil dilation velocity (Fig. 4D; Exp 2B) may reflect the recovery from the insensitivity during the period of constant perturbation direction between the 8th and 9th change points (Fig. 3B), although the interaction between change point (8 and 9) and group (Exp 2A and 2B) remained marginal (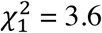, *p* = 0.057, likelihood ratio test between models with vs. without group × change point interaction).

Given the decline in pupil responses to repeated change points, these results, including those of Exp 1, further indicate that both phasic and tonic pupil responses are not only driven by errors, which was evident in the responses to the first change point, but are also likely to be modulated by factors other than fatigue or boredom. These factors may include novelty or knowledge about task structure (e.g., the presence of change points, or type/direction of perturbation), which would presumably make large errors less surprising. When we looked closely at the kinematic errors at the change point trials (i.e., the exact trial of force introduction/reversals), some evidence indicated that participants gradually became better at handling sudden changes in mechanical perturbations. The size of trajectory errors at the point of peak velocity (PVE) for the change point trial #2 through #10 (i.e., unexpected reversal in force field) exhibited a systematic reduction as participants experienced more reversals (Supplementary Figure 2A; *t*_177_ = −3.91, *p* = 1.31 × 10^−4^, a significant negative slope for change point number assessed by a linear mixed-effects model). Such a reduction in PVE was not simply caused by a similar reduction of peak y-velocity in these change point trials (tendency towards increase: *t*_175_ = 1.94, *p* = 0.054, slope assessed by a linear mixed-effects model; Supplementary Figure 2C), or a systematic reduction in learning in feed-forward motor commands in the previous force field environment (no systematic decline in learning index measured at the closest channel trial before each change point: *t*_193_ = −0.43, *p* = 0.67, linear mixed-effects model; Supplementary Figure 2B). Note that, as already mentioned, the decrease in pupil responses was significant even after taking this reduction of kinematic errors into account.

### Pupil responses to gradual force perturbation: Association between baseline pupil diameter and perturbation awareness

We also ran another experiment with 34 new participants in which they adapted to a gradually introduced force field. The magnitude of the force field was gradually increased, as shown in Figure 5A (see Methods for details). As previously noted, the gradual introduction of perturbations is believed to evoke substantially less awareness about the presence of perturbation compared with abruptly-introduced perturbations because of the smaller errors it induces (50, 51). As expected, the magnitude of error experienced by participants was much smaller compared with Exp 1 and 2 (Fig. 5B). As shown in Figure 5C, participants gradually adapted to the force field to a level that was comparable to that shown by participants in Exp 2C (Supplementary Figure 1B). There was no significant difference in learning indices between the average of the last five channel trials in the CW field for Exp 2C vs. the average of the last five channel trials before the start of constant channel phase for Exp 3 (*t*_41_ = −1.04, *p* = 0.31, two-sided independent samples *t*-test). How does the pupil respond to gradual perturbation?

**Figure 5.**
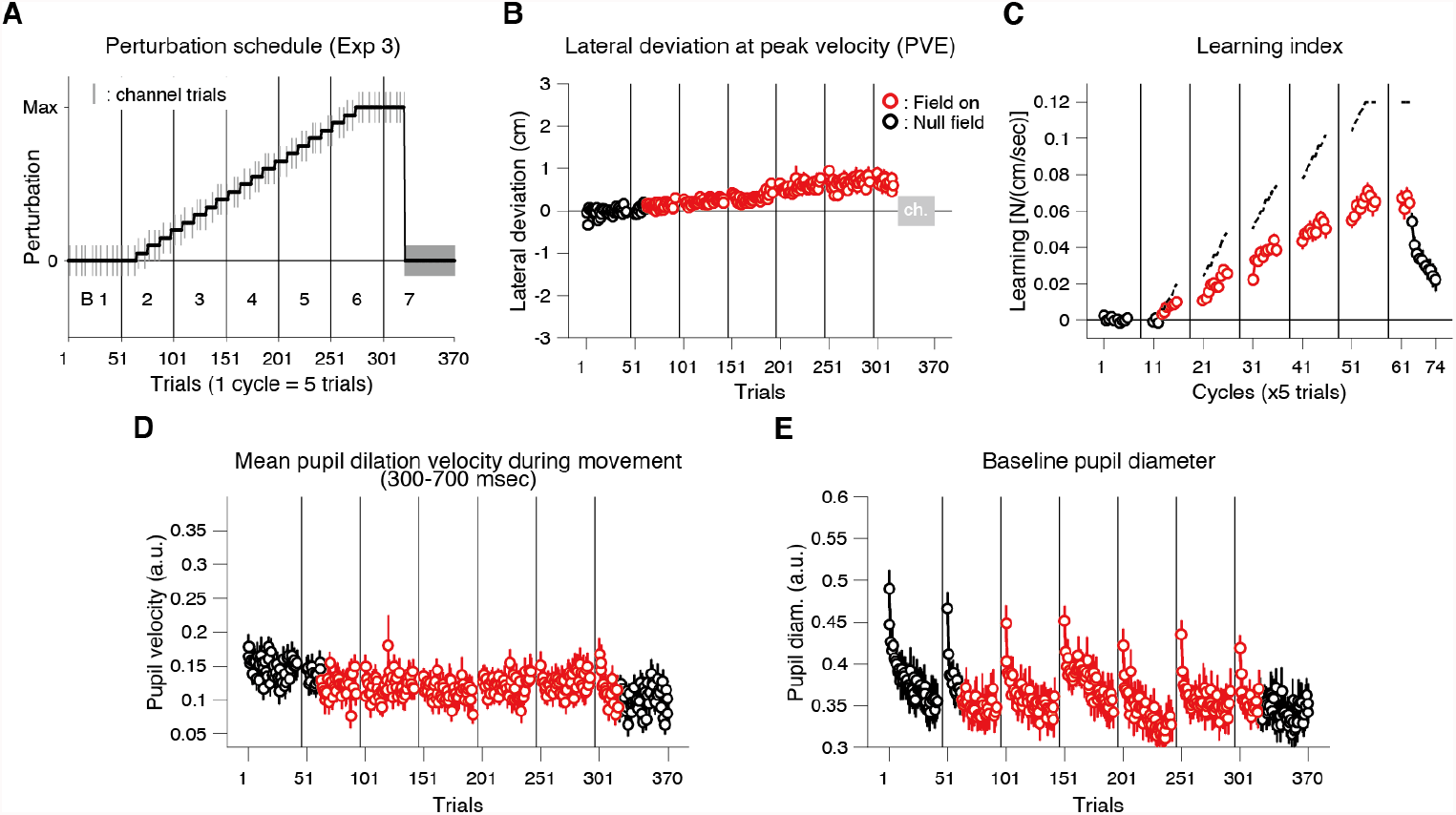
Pupil responses to gradually introduced force field (Experiment 3). **(A)** The perturbation schedule for Exp 3 (n = 34). The size of the force field was gradually increased with small steps (see Methods). The vertical lines indicate set breaks. The short gray vertical lines represent the force channel trials. **(B)** Mean lateral hand deviation at the peak tangential handle velocity for each trial. Positive values correspond to rightward deviation. **(C)** Mean learning index measured in the channel trials (once in a cycle). Learning index was the lateral force to the channel at the time of peak velocity divided by the peak velocity (i.e., viscosity). Dashed lines indicate ideal values for the learning index (panel A) averaged within each cycle. **(D)** Mean pupil dilation velocity (averaged between 300 to 700 ms since the movement onset) for each trial. **(E)** The baseline pupil diameter for each trial. The red dots represent values for perturbation trials (B–E). Error bars correspond to s.e.m. across participants (B–E).

As expected, except for the clear and consistent increase in baseline pupil diameter at the start of new block, participants’ pupils showed no clear responses, in terms of both phasic and tonic activity (Fig. 5D, E). The results clearly indicate that the pupil responses were more sensitive to a sudden and large change in the environment (Exp 1 and 2) than to a covert, gradual change (Exp 3), suggesting some similarity between participants’ awareness about the perturbation and subjective uncertainty about the task state change reflected in baseline pupil diameter.

To further investigate the relationship between pupil responses and awareness about perturbation, we assessed participants’ awareness of perturbation immediately after the experiment by asking them to report the presence/absence of force perturbation in each block without providing any information about this interview prior to the main session (see Methods for more detail). Based on these data, we split participants into two groups according to the total number of blocks in which they reported that perturbation was present. Figure 6A shows the resultant median-split report rate and their average. First, the data showed a gradual increase in report rate, consistent with the gradual increase in the force field. Interestingly, although the perturbation was gradually introduced, on average, more than half of participants were aware of its presence in the later stage of learning, in which the perturbation size almost reached its maximal value (blocks 5 and 6). The chi-squared test revealed a significant difference in %presence between the sub-groups in blocks #4 (*𝓍*^2^ = 6.47, *p* = 0.01), 5 (*𝓍*^2^ = 11.52, *p* = 6.896 × 10^−4^), and 6 (*𝓍*^2^ = 9.49, *p* = 0.002). There was no significant bias in applied force direction between the sub-groups (*𝓍*^2^ = 0.008; *p* = 0.9285), indicating that the difference was not simply caused by the different sensitivity to force direction.

**Figure 6.**
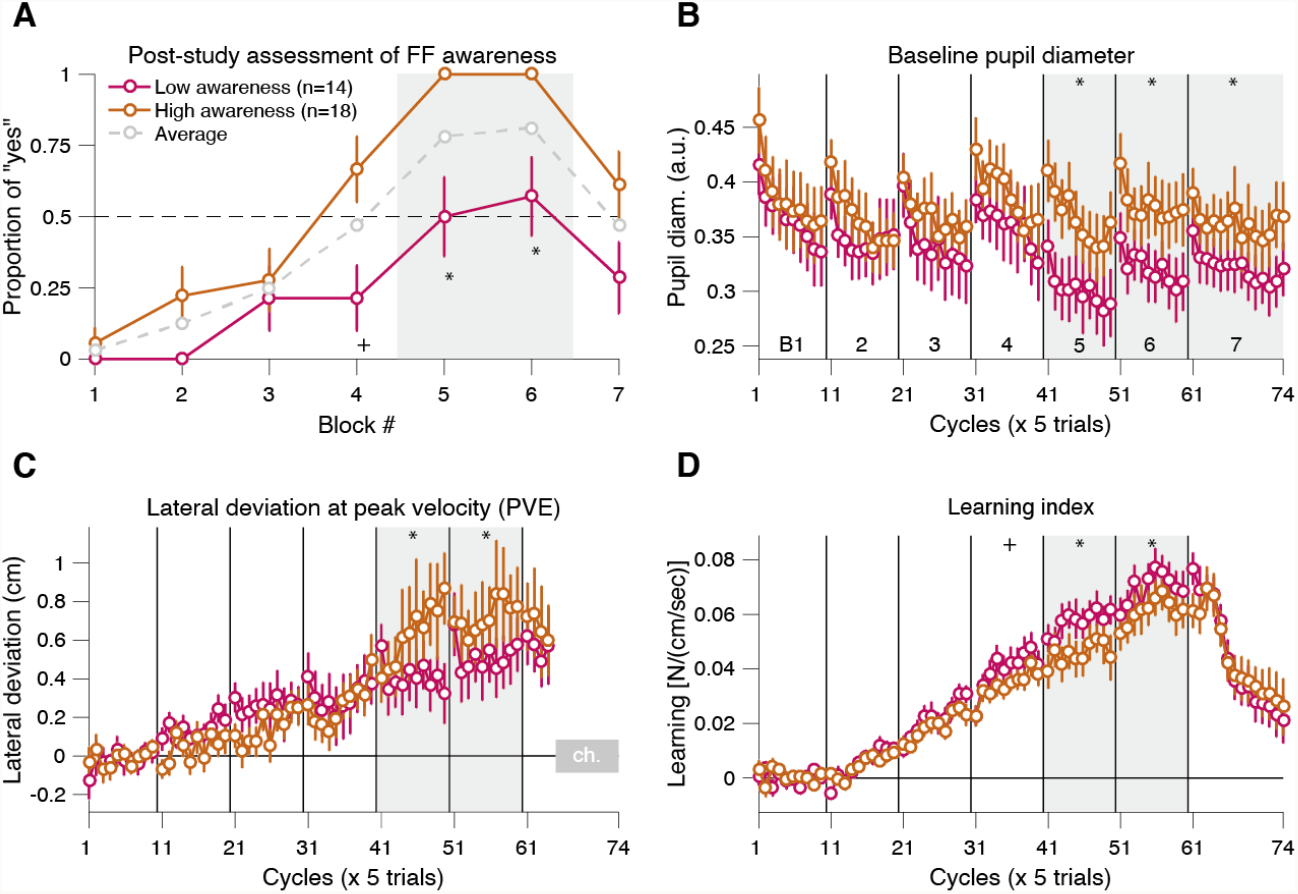
Individual differences in baseline pupil diameter were correlated with awareness of the force field (Experiment 3). **(A)** The score of the post-study questionnaire for awareness of the force field perturbation for each block. Data were split into two sub-groups according to the median of the total score (number of “yes” responses) over the blocks (below-median: purple dots, above-median: gold dots). Gray dots indicate the mean score for all participants. **(B–D)** Median split data for baseline pupil diameter (B), PVE (C), and learning index (D). Data were averaged within each cycle (five-trial bins). Error bars correspond to s.e.m. across the participants. Gray shaded areas indicate a significant difference (chi-squared test for A; linear mixed-effects model for B–D). Asterisks (*) indicate a significant effect after correcting for multiple tests using the Holm-Bonferroni method controlling for family-wise error rate (*p* = 0.05). Plus (+) signs indicate significance assessed using the Benjamini-Hochberg method controlling for false discovery rate (*q* = 0.05). Raw *p*-values are reported in the main text.

Intriguingly, comparison of pupil data between sub-groups revealed that baseline pupil diameter was larger for the “more aware” participants in blocks 5 (*F*_1,2286.5_ = 26.57, *p* = 2.75 × 10^−7^), 6 (*F*_1,2286.5_ = 19.25, *p* = 1.20 × 10^−5^), and 7 (*F*_1,2286.5_ = 7.48, *p* = 0.006), detected as significant sub-group × block interaction terms assessed by a linear mixed-effects model (see Methods for more detail). This was accompanied by larger movement error (maximally ∼0.2 cm) in blocks 5 (*F*_1,1971.5_ = 10.64, *p* = 0.001) and 6 (*F*_1,1971.5_ = 8.62, *p* = 0.003) and smaller learning index values in blocks 4 (*F*_1,2286.1_ = 5.31, *p* = 0.02), 5 (*F*_1,2286.1_ = 19.75, *p* = 9.24×10^−6^), and 6 (*F*_1,2286.1_ = 9.79, *p* = 0.0017) in the “more aware” participants. These results suggest that baseline pupil diameter is likely to be associated with participants’ awareness about the perturbation. This result is also consistent with the notion that the increase in tonic baseline pupil diameter reflects increased subjective uncertainty about the environment.

### Increased baseline pupil diameter at the start of new blocks suggests increased subjective uncertainty

The most prominent feature for the pupil responses in Exp 3 was a characteristic re-increase in baseline pupil diameter at the beginning of a new block (Fig. 5E). This phenomenon was robustly accompanied by extended reaction time (RT) and movement time (MT). Interestingly, these features were also consistently observed in Exp 1 and 2. Analyzing the data from all of the experiments revealed a consistent pattern of modulation between baseline pupil diameter, RT and MT, all of which showed larger values in earlier trials in the block (see Method for detail). The comparison between the average of early vs. late trials consistently yielded significant differences between them (statistics summarized in the Table 2). Notably, if the increased tonic baseline pupil diameter at the start of the new block merely indicated higher arousal/vigilance/vigor because of short rests between the blocks, the RT and MT would be expected to decrease (26, 27, 52). Longer RTs typically accompany decisions under uncertain condition or choices with many potential options (Hick’s law), and longer MTs are observed when the task is perceived as difficult (Fitts’s law). Therefore, such increases in baseline pupil diameter should, at least in part, reflect an internal state related to increased subjective uncertainty.

**Table 2.** Statistical values for the comparison between early vs. late trials after set breaks (Fig. 7).

| Experiments | Variables | *t*-values | df | *p*-values |
| --- | --- | --- | --- | --- |
| Exp 1 | Baseline pupil diameter | 3.06 | 27 | 0.005 |
|  | Mean pupil dilation velocity | 1.83 | 27 | 0.078 |
| | Reaction time | 4.27 | 27 | $2.13 \times 10^{-4}$ |
| | Movement time | 6.34 | 27 | $8.58 \times 10^{-7}$ |
| Exp 2 | Baseline pupil diameter | 8.99 | 28 | $9.57 \times 10^{-10}$ |
|  | Mean pupil dilation velocity | 2.95 | 28 | 0.006 |
| | Reaction time | 4.70 | 28 | $6.23 \times 10^{-5}$ |
|  | Movement time | 3.41 | 28 | 0.002 |
| Exp 3 | Baseline pupil diameter | 9.26 | 33 | $1.07 \times 10^{-10}$ |
|  | Mean pupil dilation velocity | 2.34 | 33 | 0.026 |
| | Reaction time | 6.29 | 33 | $4.13 \times 10^{-7}$ |
| | Movement time | 4.88 | 33 | $2.64 \times 10^{-5}$ |

Two-sided paired *t*-tests were used for the comparison.

## Discussion

In the current study, we systematically studied pupillary responses during the process of force field reach adaptation in which the perturbation was applied either abruptly (Exp 1), abruptly with multiple reversals of the force direction (Exp 2), or gradually (Exp 3). The results can be summarized as follows: (i) unexpectedly large movement error led to increased phasic pupil dilation during movement that was accompanied by a transient increase in tonic baseline pupil diameter in subsequent trials (Exp 1–3); (ii) such pupil responses to error were attenuated when participants repeatedly experienced large errors induced by multiple reversals (Exp 2); (iii) baseline pupil diameter showed consistently larger values at the beginning of each experimental block, which were consistently accompanied by increased reaction time and movement time (Exp 1–3); and (iv) post-study awareness of gradually-introduced force field correlated with baseline pupil diameter during the task (Exp 3). These results constitute the first detailed characterization of pupillary responses during human motor learning.

As discussed below, our results support an interpretation that is generally consistent with proposals in the field of cognitive, reward-based learning (12–18, 30, 28, 29), suggesting that baseline pupil diameter is likely to reflect subjective uncertainty about the environment, whereas phasic pupil dilation during movement is likely to reflect surprise about sensory consequences (e.g., movement error), and both are modulated by the novelty of the environment. At the beginning of a new block, when the novelty of a task is presumably high because of participants’ lack of complete knowledge about the experiment or forgetting during the break, both the tonic baseline pupil diameter and phasic pupil dilation were high. In contrast, at change points in which the environment (i.e., force field direction) is switching between known conditions, at which time the novelty of an error is low, pupil responses are less sensitive to the size of the observed error. These processes may, at least in part, be mediated by central noradrenaline (NA) activity.

### What do pupil responses reflect during motor adaptation?

Neurophysiologically, pupil diameter under constant luminance is known to reflect the activity of the locus coeruleus (LC) in the brainstem, a central source of NA (53–55). As suggested by numerous studies, pupil size and LC neurons show similar response patterns to surprising events and perceived environmental uncertainty (53, 54). For instance, both LC neurons and the pupil show phasic responses to novel, infrequent, or surprising stimuli (60, 56, 18, 57–59, 14, 12, 13, 61, 15–17). Moreover, LC neurons and the pupil show a transient increase in tonic activity facing the reversal of target-reward contingency (i.e., sudden increases in prediction error) in both monkeys (61) and humans (12–17, 30, 28, 29). Thus, phasic pupil dilations and tonic increases in baseline pupil diameter are likely to reflect perceived surprise and environmental uncertainty, respectively, mediated by corresponding activation in the LC-NA system. Thus, the present results can be interpreted as follows. A sudden introduction/reversal of the force field induced substantial surprise, leading to a phasic response in the pupil-linked NA/arousal system. This is then followed by a transient increase in uncertainty regarding the task environment and a corresponding increase in tonic activity of the pupil-linked NA/arousal system (Exp 1, 2). The absence of such pupil responses in the gradually increasing force field condition (Exp 3) suggests that the size of (prediction) error is a key factor driving this process.

Interestingly, further observation of the results of Exp 2 indicated that pupil responses to errors might also be modulated by novelty. The decline in pupil responses to large errors after several change points (Fig. 4B, D) indicates that error size is not the only determinant of pupil responses during motor adaptation. Conceivably, acquisition of higher-level knowledge about task structure (e.g., the presence of reversal) might have made the sudden increase in movement errors no longer surprising, but somehow expected. A recent monkey study using a choice-reversal task also reported that the more reversals monkeys experienced, the faster they switched behavior, which was captured by a Bayesian choice model as gradually increasing prior belief on reversal (62). In accord with this notion, we observed a gradual reduction in kinematic error in the change point trials, which was not attributable to reduced learning in the preceding force direction or reduced peak hand velocity (Supplementary Figure 2). These results indicated that participants became gradually better at moving under the force field, including sudden reversals, potentially because of increased stiffness (63, 64), improved feedback control (65, 66), or both. It should be noted that the decline in pupil sensitivity to frequent increases in errors, as observed in the present study, has not been described for the similar reversal learning paradigm in the context of reward-based learning (e.g., 14, 15). Although it is not yet clear whether this phenomenon is specific to motor learning, our results extend the findings of previous studies in revealing how the pupil-linked arousal/NA system responds to changes in the environment.

We speculate that the larger baseline pupil diameter observed at the beginning of new experimental blocks reflects heightened subjective uncertainty about the task (e.g., Fig. 5E; Exp 3). This characteristic pattern of larger pupil size early in experimental blocks has also been frequently reported but often interpreted simply as (re-)increase in arousal after short rests (e.g., 68, 69). However, as indicated by the increased RT and MT in this phase (Fig. 7), the larger pupil diameter here was more likely to be associated with increased subjective uncertainty because RT typically increases when facing uncertain decisions (69) or choices with many potential options (70), and MT typically increases when the task is perceived as difficult (i.e., subjective expectation of movement accuracy is low) (71). A recent study also reported a larger pupil diameter and slower RT for uncertain decisions (72). If larger baseline pupil diameter only reflects higher arousal/wakefulness (22–25), vigilance/alertness (26, 27), and/or vigor (52, 73), RT and MT would be expected to decrease (Fig. 7D). Thus, our data suggest that larger pupil diameter after set breaks at least partially reflects subjective environmental uncertainty.

**Figure 7.**
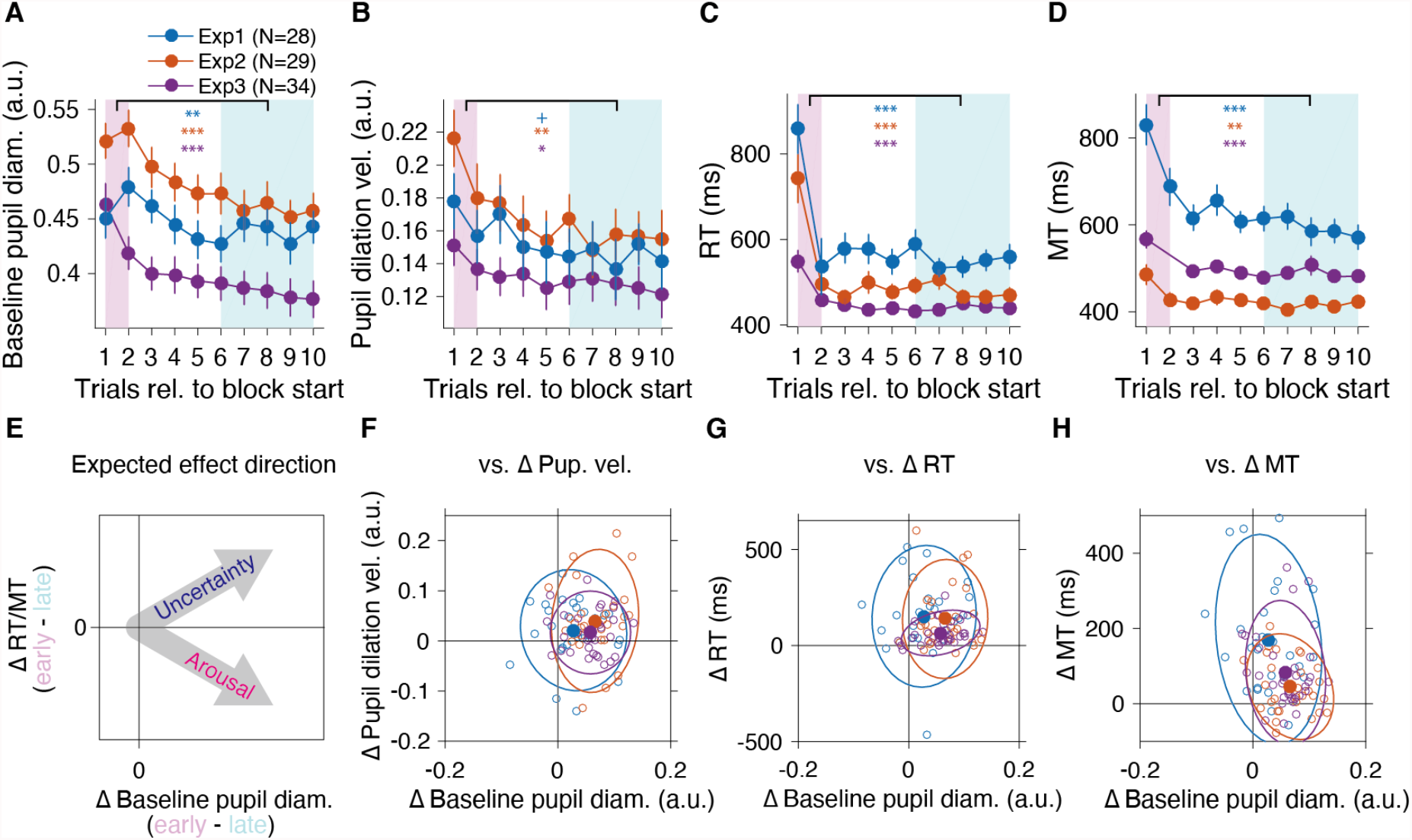
Dilated baseline pupil diameter at the start of a block implies increased subjective uncertainty (Experiment 1–3). **(A–D)** Change in baseline pupil diameter (A), pupil dilation velocity (B), reaction time (C), and movement time (D) for the first 10 trials after the set break, averaged over multiple blocks (see Methods). Error bars indicate s.e.m. across participants. **(E)** A schematic illustration of the association between the delta-baseline pupil diameter and delta-RT or -MT expected either from uncertainty or an arousal/vigilance account of baseline pupil diameter (arrows). **(F–H)** Scatter plot for delta-baseline pupil diameter vs. delta-pupil dilation velocity (F), delta-RT (E), or delta-MT (F), calculated as the difference between averages indicated by colored areas (pink: early trials, cyan: late trials) in panels A–D. In all cases (F, G, and H), the group-level effect lies in the first quadrant, consistent with the uncertainty account of increased baseline pupil diameter illustrated in panel E. Solid closed dots represent averaged data. Open dots represent individual data. Ellipses indicate the 95% confidence contour. Asterisks (colors correspond to experiment as indicated in panel A) indicate significance assessed by two-sided paired *t*-test between the averaged data for early vs. late trials (+: *p* < 0.1, *: *p* < 0.05, **: *p* < 0.01, ***: *p* < 0.001).

### How can pupil-linked arousal influence motor adaptation?

The transient increase in subjective uncertainty in the environment and pupil-linked arousal/NA activity induced by the sudden introduction of force fields bears a similarity to several phenomena reported in the early phase of human motor adaptation. For example, adaptation to both stable and unstable force fields transiently increases muscle co-activation (74, 64, 75), as increasing limb impedance is an optimal strategy for movement under highly uncertain dynamic environments (63) or to increase movement accuracy (76). Similarly, adaptation to a force field transiently increases the gains of visuomotor feedback responses (77), as well as long-latency muscle stretch reflex (78), which is mediated by the primary motor cortex (79–81). Furthermore, a recent study demonstrated that the Ia afferent firing from muscle spindles is enhanced in the early phase of visuomotor adaptation (82). These central and peripheral gain control changes might originate from the NA projections to the motor cortices and spinal cord (83). Notably, a recent report showed an arousal-like transient increase in neuronal population activity in the primary motor cortex in response to errors caused by environmental (brain-computer interface mapping) change, which correlated with pupil diameter change (68). Additional evidence suggests increased muscle spindle sensitivity following sympathetic up-regulation (84, 85).

One intriguing question is whether/how pupil responses can be informative for understanding the “explicit” and “implicit” components of motor learning (86, 5, 6). Currently, the dominant approach for trial-by-trial assessment of the conscious/explicit component of motor learning in the reach adaptation paradigm is to ask participants to verbally/manually report their aiming direction prior to each reach (87, 5, 88). However, this approach suffers from the inherent problem of interfering in the learning system itself and hence biasing learning processes to be more “explicit” (89–91). Moreover, although this aim reporting method allows researchers to assay the contribution of explicit processes to net learning, it does not provide direct clues regarding the underlying cognitive processes driving the explicit process. Thus, it is important to develop new complementary approaches to assess the explicit and/or implicit component (e.g., 92, 93) or to examine cognitive states that can drive it without affecting the learning system. Given the notion that motor learning is learning about movement selection and execution (1) and the significant effect of surprise/uncertainty (and pupil-linked arousal/NA system) on reward-based learning of action selection (15), pupil responses may provide a window for assessing how learning about movement selection (i.e., explicit process) progresses.

Although we did not explicitly quantify the explicit components, our results may provide some insight into the question above. The post-experiment questionnaire in Exp 3 (gradual force field) revealed the association between the inter-individual differences in the overall correctness of perturbation awareness and baseline pupil diameter (Fig. 6B). Such inter-individual differences were also accompanied by the amount of learning and movement error, in a manner in which higher awareness was associated with less learning and more error (Fig. 6C, D). One possibility is that participants with weaker adaptation experienced more error, resulting in larger baseline pupil diameter and an increased probability of later recall of perturbation awareness. Recent evidence also suggests that prediction error (94, 95), as well as phasic pupil responses (96, 97), can signal a subjective belief about environmental change, which helps to create event boundaries in a memory structure (96, 95). Another possibility is that participants with larger baseline pupil diameter had higher subjective uncertainty about the task (and were hence more likely to recall it as “perturbed” afterward), leading to more frequent exploratory behavior, which resulted in a smaller amount of adaptation. This scenario is consistent with previous reports that the awareness of perturbation reduced the amount of implicit adaptation (51, 98, 99) and a more recent study suggesting that the implicit adaptation process compensates for the noisy explicit strategy (88). One caveat is that, although we instructed our participants to aim straight to the target, we did not directly measure the explicit component of adaptation. Thus, at this point, we can only suggest that the tonic baseline pupil diameter reflects subjective uncertainty about the environment and may also be related to awareness about the perturbation. Further confirmatory studies will be needed to clarify this point.

### Limitations and open questions

Despite the established link between pupil diameter and central LC-NA activity, the relationship is not necessarily one-to-one. For instance, a recent rodent study, in which cortical axons for both NA and acetylcholine (ACh) were recorded, reported that while rapid changes in pupil diameter and its time derivative (i.e., dilation velocity) were more strongly correlated with NA than ACh activity, slow pupil dilations on the timescale of a few seconds were correlated with activation of both NA and ACh (100). However, the detailed mechanisms by which central ACh affects pupil size remain unknown. Moreover, recent studies have reported that the activity of the dorsal raphe nucleus (DRN) serotonin (5-HT) neurons in mice also tracks environmental uncertainty (101) and that photoactivation of DRN 5-HT neurons also elicits pupil dilation (102). However, the bidirectional connection between DRN and LC (83) further complicates this relationship. It is also important to note that 5-HT plays a key role in controlling the input-output gain of spinal motoneurons (103, 104). Overall, a more direct approach, such as invasive animal studies or pharmacological manipulation, is required to further establish the links among pupil diameter, these neuromodulators (NA, ACh, and 5-HT), and motor learning processes.

One topic that remains to be addressed is the relationship between motor learning rate and uncertainty/surprise and the pupil-linked arousal/NA system. Theoretically, statistically-optimal learning algorithms take multiple sources of uncertainty into account to dynamically modulate the learning rate, such as the Kalman filter (105) and more recent extensions (33, 32, 34, 106–108). Unfortunately, it was not easy to directly answer this question with the current data, because the current experiments were not designed for the accurate quantification of learning rate. One recent study (67), however, reported an association between baseline pupil diameter and learning rate in a modified saccade adaptation task. We will address this question in a separate report in which we directly measure the single-trial learning rate with/without experimental manipulation of the pupil-linked arousal system in the motor adaptation paradigm.

In the present study, we provided the first detailed characterization of pupillary responses in a widely used motor adaptation paradigm. Our data revealed how the internal states of human participants, most likely surprise and uncertainty about the environment, dynamically change during motor adaptation, thus providing important clues for understanding the process of force field learning. The results of the current study highlight the utility of pupil diameter as a valuable window into the motor system.

## Acknowledgements

We thank J. Diedrichsen, S. Kitazawa, and J. A. Pruszynski for comments on the early version of the manuscript. We also thank N. Hagura, M. Hirashima, and T. Ikegami for helpful discussions and comments on the manuscript, and S. Kato for assistance in data collection. The present work is supported by JSPS Fellowships for Young Scientists (#15J03233) and KAKENHI (#18K17916) to A.Y., and post-doctoral fellowships from NSERC and the BrainsCAN program at Western University to J.W.

## Author contributions

Atsushi Yokoi: Conceptualization, Data Curation, Formal Analysis, Funding Acquisition, Investigation, Methodology, Project Administration, Resources, Software, Supervision, Validation, Visualization, Writing – Original Draft, Writing – Review & Editing Jeffrey Weiler: Investigation, Funding Acquisition, Writing – Review & Editing

## Supplementary Figures

**Supplementary Figure 1.**
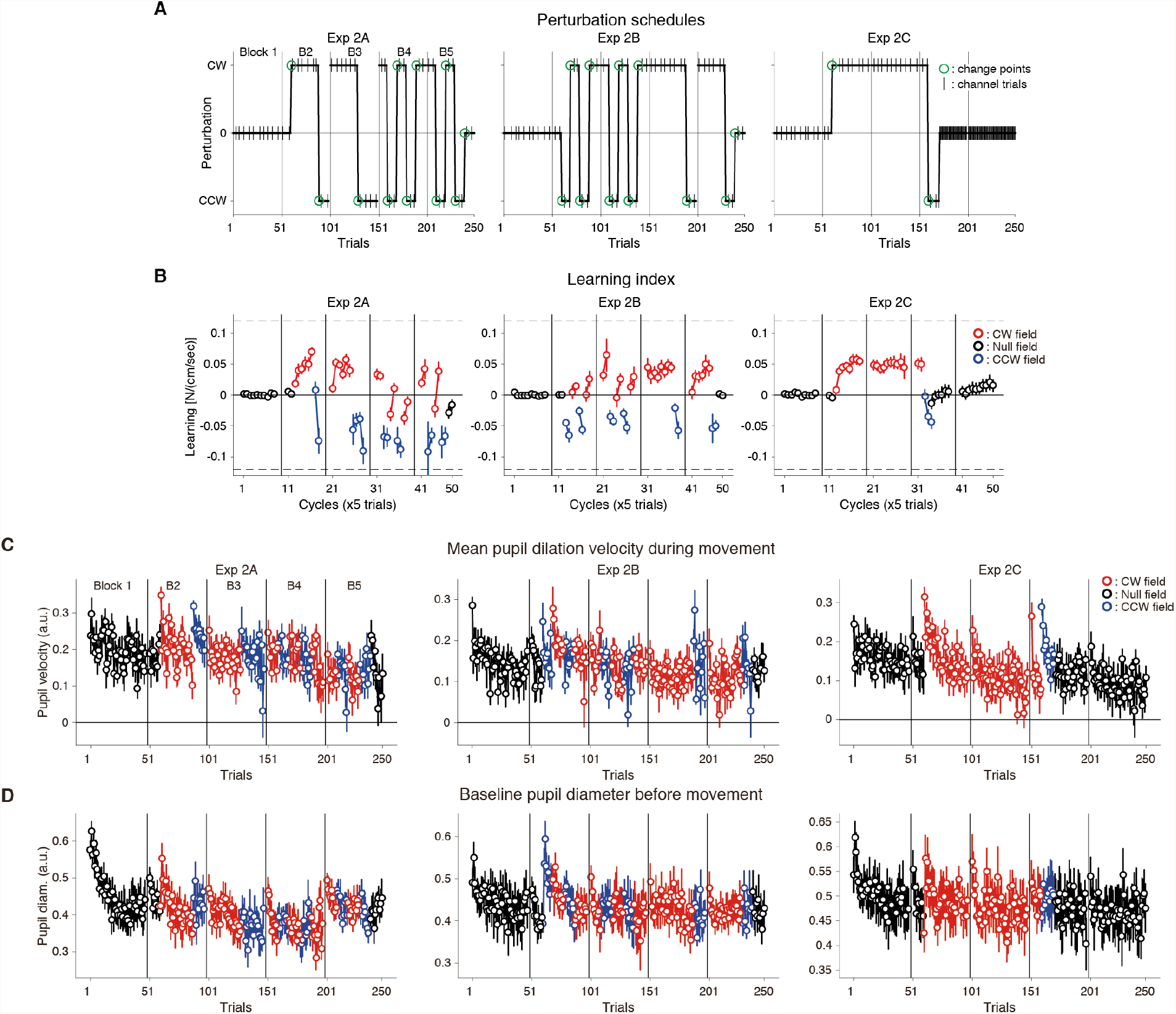
Learning index and pupil responses for Exp 2. **(A)** Perturbation schedules for the Exp 2 [Exp 2A (n=10), 2B (n=10), and 2C (n=9)]. The green circles indicate the ‘change point’ trials where either magnitude (on/off) or direction (CW/CCW) of the force field changes in the middle of the blocks (changes across the blocks are excluded). The vertical lines indicate set breaks. **(B)** Averaged learning index measured in the channel trials (once in a cycle) for the Exp 2A (first column), 2B (second column), and 2C (last column), respectively. Learning index was the lateral force to the channel at the time of peak velocity divided by the peak velocity (i.e., viscosity). **(C, D)** Trial-by-trial change in the evoked pupil dilation velocity (C), and baseline pupil diameter (D) for the Exp 2A (first column), 2B (second column), and 2C (last column), respectively. The colors of circles (black, red, and blue) indicate the data for baseline, CW field, and CCW field, respectively. Positive values for panel D correspond to rightward deviation. The vertical lines indicate set breaks. Error bars represent s.e.m. across the participants. Trials at which the dot color changed indicate the change points.

**Supplementary Figure 2.**
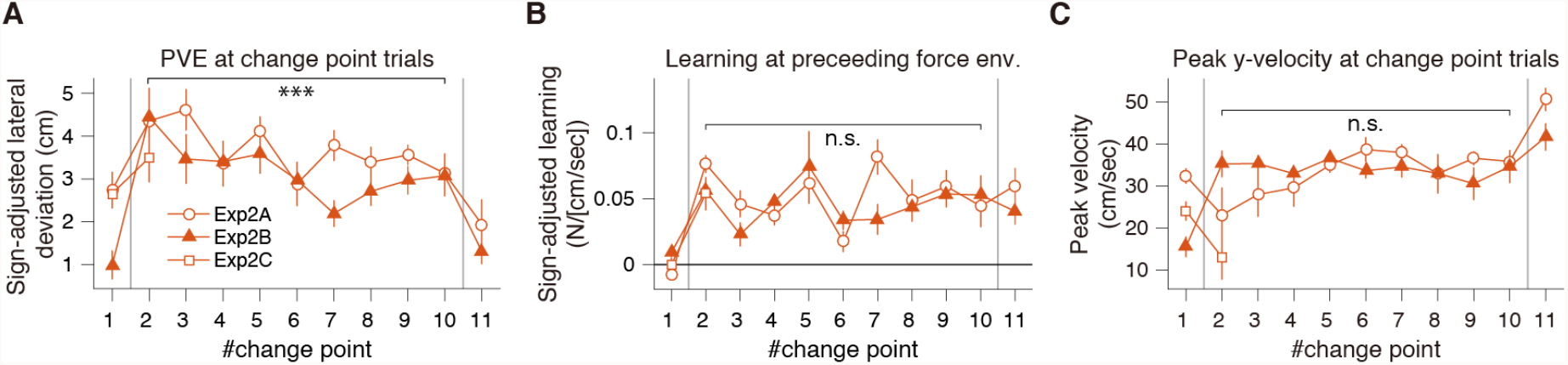
Change in kinematic errors over multiple change points (Exp 2). **(A)** Sign-adjusted PVE at each change point (the first trial of force introduction/reversals) showed a gradual reduction from the second through the tenth change point, after the increase due to reversal in force direction from the first to the second. **(B)** Sign-adjusted learning index (lateral force at peak velocity divided by the peak velocity) measured at channel trials immediately before each change point (e.g., value for the second change point indicates the last measured learning index in the first force field period). Although there were some fluctuations, there was no systematic reduction. **(C)** Peak hand y-velocity at each change point. Symbols (*** or n.s.) indicate the significance of slope value in terms of #change point estimated by linear mixed-effects model from the second through the 10th change points indicated by gray vertical lines (n.s.: not significant; ***: *p*<0.001). The data for Exp 2C was not included for the linear mixed-effects model fitting as there were only two change points in this sub-group.

